# *Toxoplasma gondii* bradyzoites induce transcriptional changes to host cells and prevent IFNγ-mediated cell death

**DOI:** 10.1101/669689

**Authors:** Simona Seizova, Alexandra L Garnham, Michael J Coffey, Lachlan W Whitehead, Kelly L Rogers, Christopher J Tonkin

## Abstract

*Toxoplasma gondii*, the causative agent of toxoplasmosis, lies dormant for life and is a reservoir for disease reactivation, causing blindness, encephalitis and congenital birth defects. Acute-stage tachyzoites extensively manipulate their host cell by exporting a repertoire of proteins across the parasitophorous vacuolar membrane (PVM). This interferes with the hosts transcriptional program, allowing for persistence during immune attack. It is unknown how bradyzoites persist and what role host manipulation plays in latency. Here we show that bradyzoite-containing host cells have a unique transcriptional landscape when compared to tachyzoite infection. We demonstrate that many of these changes are dependent parasite protein export. Furthermore, we show that bradyzoite effector proteins protect host cell’s from IFNγ-mediated cell death, thus highlighting the functional importance of host manipulation. Together, our work provides the first understanding of how *Toxoplasma* sets up latency to persist in its host.

## Introduction

*Toxoplasma gondii* is an obligate intracellular parasite that can infect many warm-blood animals causing toxoplasmosis. *Toxoplasma* infects one third of the human population and is contracted by eating undercooked meat that harbour the cysts forms or consumption of food contaminated with oocysts shed in cat faeces^1^. Acute toxoplasmosis results in a life-long chronic infection, whereby bradyzoites establish tissues cysts in the central nervous system and muscle tissue (Dubey et al., 1998). Chronic infection in healthy individuals is largely asymptomatic; however, severe complications can arise in immune-compromised individuals from the reactivation of encysted bradyzoites. Furthermore, reactivation of tissue cysts in retinal tissue is a major cause of blindness in some countries^2, 3^. In recent years, chronic infection has also been associated with a several psychiatric disorders, suggesting that latent *Toxoplasma* may subtly effect the brain in unappreciated ways^4^. There are currently no treatments to clear the bradyzoite reservoir to prevent reactivation and associated diseases, or an understanding of how latent forms manipulate the brain to survive and cause alterations to brain physiology.

Like many pathogens, *Toxoplasma’s* ability to survive within its host can be attributed to its capacity to avoid immune detection^5^, achieved, in part, by their ability to export parasite effector proteins that interfere with host cell programs. Acute stage tachyzoites are able to modulate the host using two distinct temporally separated pathways. The first wave of effectors is injected into the host cell during the invasion process, whilst the second occurs during intracellular growth^6^. During invasion, tachyzoites inject rhoptry (ROP) proteins into the host cell, which can traffic to the outer side of the parasitophorous vacuolar membrane (PVM)^7^ and host cell nucleus^8^. Characterised ROP effectors act by interfering with the immune response by either altering host gene expression^9^ by directly disrupt signalling pathways (e.g. STAT3 and STAT6)^8, 10, 11^ or preventing destruction of the vacuole by interfering with function of key host proteins induced by IFNγ signalling (e.g. IRGs and GBPs)^7, 12, 13^.

After invasion is completed, tachyzoites activate a second wave of protein export by secreting proteins from the parasite’s dense granule (DG; DG proteins annotated by GRA) organelles. Some of these proteins sit at the host interface, often interacting with host proteins in order to modulate their activity^14–16^. For example, GRA15 localises to the host cytosolic face of PV/PVM, where it manipulates NFκB signalling, initiating IL-12 secretion and interferon gamma (IFNγ) production^14^, whereas MAF-1 protrudes from the vacuolar membrane and sequesters the host mitochondria to surround the PVM^15^. Additionally, a subset of dense granule proteins are translocated across the PVM into the cytosol or nucleus of the host. These include cell cycle regulator GRA16^17^, p38α MAPK activator GRA24^18^, Inhibitor of <u>S</u>TAT1 <u>T</u>ranscription (IST)^19, 20^, GRA28 (effects on host are unknown)^21^, β-catenin signalling modulator GRA18 is associated with anti-inflammatory responses^22^ and, more recently, HCE1/ TEEGR, which has been shown to affect the cyclin proteins^23^ and host immune regulation by the suppression of NF-κB-regulated cytokines^24^. Interestingly, all the exported dense granule proteins are predicted to be disordered, which could be essential to the translocation process^25^.

All known effector proteins traffic via *Toxoplasma’s* secretory pathway^26^ and are translocated across the PVM using parasite-derived machinery^25, 27, 28^. Upon trafficking through the parasite secretory pathway, some effectors are matured by a resident Golgi Aspartyl Protease, ASP5. ASP5 cleaves proteins at the TEXEL (*<u>T</u>oxoplasma <u>Ex</u>port <u>El</u>ement*) ‘RRL’ motif, and is required for PVM translocation of all tested DG effector proteins^19, 22, 27, 29, 30^. This has similarities with the export process of proteins in *Plasmodium* spp.^31–34^; however, unlike *Plasmodium* spp., it appears that not all TEXEL-containing protein are exported^18, 27, 35^ Recent evidence suggests a protein complex, consisting in part of MYR1, MYR2 and MYR3, is required for the translocation of proteins across the PVM^25, 28^, but the exact mechanism it yet to be defined. Furthermore, MYR1-dependent transcriptional changes account for a large portion of the changes in host transcription induced by tachyzoite infection, consistently affecting immune response pathways^36^

Despite its extensive ability to shut down defence mechanisms, *Toxoplasma* is tightly controlled by the innate and adaptive immune responses. IFNγ is vital in controlling acute *Toxoplasma* infection and prevents reactivation of latent bradyzoites^37^. Previous studies have used murine cell lines to understand how IFNγ can induce tachyzoite clearance. It has been shown that IFNγ stimulated guanylate-binding proteins (GBPs)^38, 39^ and immune related GTPases (IRGs)^40^ are recruited to the tachyzoite PVM, but the downstream clearance mechanism that follows remains elusive^41^. *Toxoplasma* has developed mechanisms to prevent IFNγ activation of infected cells^9, 42, 43^. One mechanism is DG protein IST, which is exported via the MYR1 pathway where it inhibits IFNγ induced STAT1 signalling and transcription by modifying the GAS (Gamma Activated Sequence) sites through the recruitment of the Mi-2/NuRD chromatin remodelling complex^19, 20^, thus blunting transcriptional changes when the host cell encounters this cytokine. It is not known if IFNγ has any capacity to affect bradyzoite clearance, even though it has consistently been shown to have an important role in immune defence *in vivo* ^44–46^.

In fact, most investigations into *Toxoplasma* host manipulation have been performed with acute stage tachyzoites, whereas bradyzoite-host cell interaction and manipulation has been largely neglected. Indeed, it is highly likely that host cell manipulation does differ between tachyzoites and bradyzoites to accommodate their different lifestyles and requirements. This must occur both when setting up a latent infection (ie. upon differentiation from tachyzoites to bradyzoites) as well as ongoing changes that must exist to allow for latent forms to persist over long time periods. Here we tackle the former to understand the differences between transcriptional changes between tachyzoites and seven-day-old bradyzoite forms. Our study shows that, at 7 days post differentiation, bradyzoite-containing host cells have a unique transcriptional landscape when compared to tachyzoite-infected cells. We demonstrate that many of these changes are imparted by both the presence of bradyzoite effector proteins, including IST, and likely the absence of tachyzoite effector proteins, which are not expressed/exported at this dormant stage. Furthermore, we show that bradyzoite effector proteins protect host cell’s from IFNγ-mediated cell death, thus highlighting the functional importance of host manipulation. Together, our work provides the first understanding of how *Toxoplasma* sets up latency to persist in its host.

## Experimental Procedures

### Host cell and parasite cultures and differentiation

All *Toxoplasma* parental parasite lines used in this study are of the ‘type II’ Prugniaud (Pru) background, either Pru-Toxofilin-Cre^47^, or Pru Δ*hxgprt* Δ*ku80* (PruΔ*ku80*)^48^. These parasites, and all subsequently derived lines, were cultured in primary HFFs (American Type Culture Collection, ATCC) in Dulbecco’s Modified Eagle medium (DME) supplemented with 1% v/v fetal calf serum (FCS) (Invitrogen, Australia) and 1% v/v Glutamax (Invitrogen) (D1). Prior to infection HFFs were grown to confluency in DME supplemented with 10% v/v cosmic calf serum (GE Healthcare, New Zealand) (D10). Cells were incubated at 37 °C with 10% CO_2_.

Bradyzoite differentiation was achieved by inoculating tachyzoites onto confluent HFFs at an MOI of 0.1 (for IFA and FACS) or 0.2 (for RNA sequencing and western blots) and cultured in RPMI-HEPES supplemented with 5% v/v fetal calf serum (FCS) (Invitrogen, Australia) at pH 8.1-8.2 and allowed to differentiate at 37 °C with ambient CO_2_ ^49^. The media was changed every 2 days (to prevent acidification) for at least 7 days prior to assays.

### DNA, plasmids and transfection

Candidate genes were targeted in all cases by the CRISPR/Cas9 system, which has been adapted for use in *Toxoplasma*^50, 51^. EuPaGDT (http://grna.ctegd.uga.edu)^52^ was used for guide selection and CRISPR-Cas9 target plasmids were made by Q5 mutagenesis (NEB) using pU6_Universal_Cas9_mCherry^27, 51^.

PruΔ*myr1* was created by transfection of *myr1* CRISPR guide and followed by serial dilution cloning and sequencing. PruΔ*myr1* knockouts were confirmed by the transient transfection of the GRA24-Myc3-expressing plasmid^27^. Candidate genes in PruΔ*ku80* were tagged with a DNA sequence encoding the HA epitope tag to allow for monitoring of protein expression during the generation of knockouts^53^. Briefly, CRISPR guide plasmids were co-transfected with either HA containing dimerised oligo primers, pLIC-3xHA-HXGPRT^35^ or, for knockouts, pLoxP-DHFR-mCherry^54^ amplicon and selected with 6-thiozanthine and mycohenolic acid, and pyrimethamine respectively (Fig. S5a)^55, 56^. 10 ug of Cas9 plasmid was combined with up to 80 ug of amplicon and resuspended in 20 uL P3 solution (Lonza), before being transfected by the Amaxa 4D Nucleofector (Lonza) using the code FI-115 (Human Unstimulated T-cells).

### IFA and antibodies

Parasites were fixed in 4% v/v paraformaldehyde in PBS for 10 min; permeabilized in 0.1% v/v Triton X-100 in PBS and blocked in 3% w/v BSA (Sigma) in PBS for 1 hr. For samples with Propidium Iodide, cells were incubated for 10mins in a final concentration of 0.05mg/ml (Merck). Samples were washed three times prior to fixation and permeabilized as described above. The following antibodies were used in this study: αGAP45^57^, αSAG1 DG52^58^, αHA 3F10 (Roche), αSRS9^59^, αCST1^59^, Fluorescein-Dolichos Biflorus Agglutinin (DBA; Vector Labs), αIRF1 D5E4 (Cell Signalling Technologies) and αMyc 9E10 (Sigma). Primary antibodies were diluted in the bovine serum albumin (BSA)/PBS solution for 1 hr, washed, and then incubated with Alexa Fluor-conjugated secondary antibodies (Invitrogen) for 1 hr. 5 μg/ml DAPI was added in the penultimate wash for 10 min and samples were mounted onto microscope slides with Vectashield (Vector Labs). Parasites were imaged using either the Zeiss AxioLive Cell Observer, or the Zeiss LSM 880 for quantification. Quantification was automated using an Image J macro developed in house (available on request). Briefly, the mean grey value (MGV) of the protein of interested was measured both within the parasite vacuole and the host nucleus. MGV for non-specific background was subtracted for all analyses before the ratio of export (nucleusMGV/vacuoleMGV) or ratio of IRF1 expression (IRF1infectedMGV/ IRF1uninfectedMGV) was calculated.

### Western Blots

Immunoblot samples were pelleted then lysed for 30 mins at 4 °C in 1% v/v Triton-X 100 (brand), 1 mM MgCl_2_ in PBS (Gibco) supplemented with final 1 x cOmplete protease inhibitors (Sigma) and 0.2 % v/v Benzonase (Merck). Samples were then combined with an equal volume of non-reducing 2x Sample buffer and 15 μl loaded onto a gel. Proteins were transferred onto nitrocellulose then blocked in 5 % w/v milk in 0.05 % Tween 20-supplemented PBS (PBS-T). Primary and secondary antibodies were diluted in milk/PBS-T. Nitrocellulose membranes were imaged on an Odyssey Fc imager (LI-COR Biosciences) using IRDye 800CW goat α-rat, IRDye 800CW goat α-mouse and IRDye 680RD goat α-rabbit antibodies. Antibodies used in this study were: αGAP45^57^, αSAG1 DG52^58^, αHA 3F10 (Roche), αSRS9^59^.

### IFN-gamma Stimulation

Human IFNγ 285-IF (R&D systems) was added from a 50 µg/µl stock solution to a final experimental concentration of 50 ng/µl. Tachyzoites were inoculated for 20 hours before a 15 hour stimulation in D1 supplemented with 25mg/ml dextran sulfate (Sigma)^60^. Bradyzoites were stimulated for 15 hours after 7 days in differentiation media.

### Compound 1 and BIPPO treatment

Compound 1 was added to a final concentration of 2µM an hour prior to the addition of IFNγ and left for indicated treatment period^55, 61^. BIPPO (5-Benzyl-3-isopropyl-1H-pyrazolo[4,3-d]pyrimidin-7(6H)-on) was added to a final concentration of 1µM and incubated for 7 mins before cells were harvested^62^.

### Live Imaging

Parasites where cultured and differentiated as described above in a 35mm polymer coverslip dish (Ibidi) for 7 days. 100ul of Propidium Iodide (1mg/ml; Merck) was added to 2mL of fresh differentiation media and either 50 ng/µl of human IFNγ 285-IF (R&D systems) was added or not. Between 15-25 cysts were tracked every 14 mins for 15 hours (overnight) on the Lecia SP8 confocal using resonant scanning.

### Flow Cytometry

Bradyzoites were differentiated as described above and treated as indicated prior to being washed with PBS and dislodged with trypsin. A minimum of 5×10^3^ cells were analysed live using a BD FACSAria Fusion cell sorter (BD Biosciences). The proportion of eGFP^+^ between unstimulated and stimulated cells was calculated. Compound 1 samples were fixed in 4% v/v paraformaldehyde/PBS before being analysed. Tachyzoite samples were treated with IFNγ as described above before being fixed in 4% v/v paraformaldehyde/PBS, permeabilized in 0.1% v/v Triton X-100/PBS and blocked in 3% w/v BSA/PBS for 1 hr. Samples were incubated in primary antibody, αGAP45^57^, overnight at 4°C, washed, and then incubated with Alexa-594-conjugated secondary antibody (Invitrogen) for 1 hr. All fixed samples were run on a BD LSR WII.

### Library Preparation and Transcriptome Sequencing

HFFs were passaged and grown in D10 media until they reached confluency. Following this, HFFs were transferred into D1 media (for tachyzoites; 24 hours) or differentiation media (for bradyzoites; 7 days) and left as uninfected (no parasites) or infected at a MOI of 0.2 with either Pru-Toxofilin-Cre (Pru (WT)) or Pru-Toxofilin-CreΔ*myr1* (PruΔ*myr1*) tachyzoites. Prior to RNA extraction, HFFs were washed with PBS, dislodged with trypsin, and sorted into infected (mCherry^+^) and uninfected (mCherry^-^) cells using the BD Influx (361) cell sorter (BD Biosciences). Total host and parasite RNA was extracted using the RNeasy kit (Qiagen). Three independent biological replicates of each condition were obtained.

An input of 100ng of total RNA was prepared and indexed separately for Illumina^TM^ sequencing using the TruSeq RNA sample Prep Kit (Illumina) as per manufacturer’s instruction. Each library was quantified using the Agilent Tapestation and the Qubit™ DNA BR assay kit for Qubit 3.0® Fluorometer (Life technologies). The indexed libraries were pooled and diluted to 1.5pM for paired end sequencing (2× 81 cycles) on a NextSeq 500 instrument using the v2 150 cycle High Output kit (Illumina) as per manufacturer’s instructions (generating 80bp paired-ends). The base calling and quality scoring were determined using Real-Time Analysis on board software v2.4.6, while the FASTQ file generation and de-multiplexing utilised bcl2fastq conversion software v2.15.0.4.

### Transcriptome analysis

The RNA sequencing reads were simultaneously aligned to the human and parasite genomes, builds hg38 and ME49 respectively, using the Rsubread aligner version 1.24.1 (tachyzoite and bradyzoite infection dataset) and version 1.27.4 (WT-Δ*myr1* bradyzoite infection dataset)^63^. For this alignment a reference was first built that contained both human and parasite genomic sequences. In all instances greater than 89% of the reads mapped to this combined genome. The number of fragments overlapping each human Entrez gene and parasite gene were summarized using FeatureCounts^64^ and NCBI RefSeq annotation. Differential expression analyses were then undertaken using the edgeR^65^ and limma^66^ software packages. The analysis of each data set was performed independently, details as follows.

### Tachyzoite and bradyzoite infected samples

For the analysis of the human count data, any gene which did not achieve a counts per million mapped reads (CPM) greater than 1 in at least 3 samples were deemed to be unexpressed and filtered from the data. Additionally, all genes without current annotation were also removed. The parasite count data (infected samples only) was filtered using edgeR’s filterByExpr function with default settings. Compositional differences between libraries for both human and parasite data were normalised using the trimmed mean of log expression ratios (TMM) method^67^. All counts were then transformed to log_2_CPM. Differential expression between the experimental groups in both instances was assessed relative to a fold-change threshold of 1.1 using linear models and empirical Bayes moderated t-statistics with a trended prior variance^68, 69^.

### WT-Δ*myr1* bradyzoite infection data

For this dataset, only the human count data was analysed. All genes which did not achieve a CPM greater than 0.7 in at least 3 samples was filtered from the data together with all genes with no current annotation. TMM normalisation was then applied to alleviate compositional differences between libraries. All counts were transformed to log_2_CPM. Differential expression between all groups was then evaluated relative to fold-change threshold of 1.1 using linear models and robust empirical bayes moderated t-statistics with a trended prior variance.

In both datasets the false discovery rate (FDR) was controlled below 5% using the Benjamini and Hochberg method. Analysis of the Hallmark gene sets from the Molecular Signatures Database was performed using limma’s kegga function. To make the heatmaps of the top 100 differential expressed genes, the expression of each gene was summarized as log_2_ reads per kilobase per million mapped reads (RPKM) with a prior count of 2. For the heatmaps of the Hallmark gene sets, gene expression was summarised as average log_2_ RPKMs. The plots themselves were created using the pheatmap software package and limma’s coolmap function.

### Statistical Analysis

Statistical analyses were conducted with Prism 7 (GraphPad). For groups of three or more, the test for significance was performed using a one-way analysis of variance (ANOVA) with the indicated post-test for correction. Groups of two were analysed using the unpaired Student’s t test and significance was determine p ≤ 0.05. The relationship between the intensity of IRF1 and the intensity of nuclear IST-HA was modelled with a non-linear curve (polynomial) using natural splines with 3 degrees of freedom using RStudio.

## Results

### Bradyzoites distinctively alter the host transcriptome

We sought to identify if bradyzoite-containing host cells had a unique transcriptional profile and whether this profile differed from tachyzoite-containing host cells. We first determined the best time point to take bradyzoite samples after inducing differentiation. We found that approximately 7 days post alkaline-induction at an MOI of 0.2^70^ was the longest period of time and density we could take the cultures without visibly compromising the health of bradyzoite-infected host cells. We then harvested corresponding tachyzoites at 24 hrs (under normal growth conditions) before infected cells were separated from uninfected ‘bystander’ cells by FACS based on the presence of mCherry (Fig. 1A and B). To determine bradyzoite purity, parasite samples were probed with anti-SRS9 antibodies and the number of positive vacuoles were counted by microscopy^59^. Quantitation demonstrated at least 90% of the population was positive for SRS9 under the differentiation conditions used (Fig. 1C (i); consistent with Fouts and Boothroyd 2007)^70^. RNA was then extracted and sequenced using Illumina chemistry on a NextSeq platform. We analysed the data for levels of BAG1^71^ and SAG1^72^, quintessential bradyzoite and tachyzoite markers, and this showed the expected enrichment and depletion in each sample respectively (Fig. 1C (ii)). We also interrogated other known stage-specific transcripts in our bradyzoites samples, which also showed the expected pattern of up- and down-regulation of genes associated with this stage (Fig. 1C (iii))^73, 74, 75^. This suggests that our preparations are highly enriched for bradyzoites, thus validating our experimental procedure.

**Figure 1.**
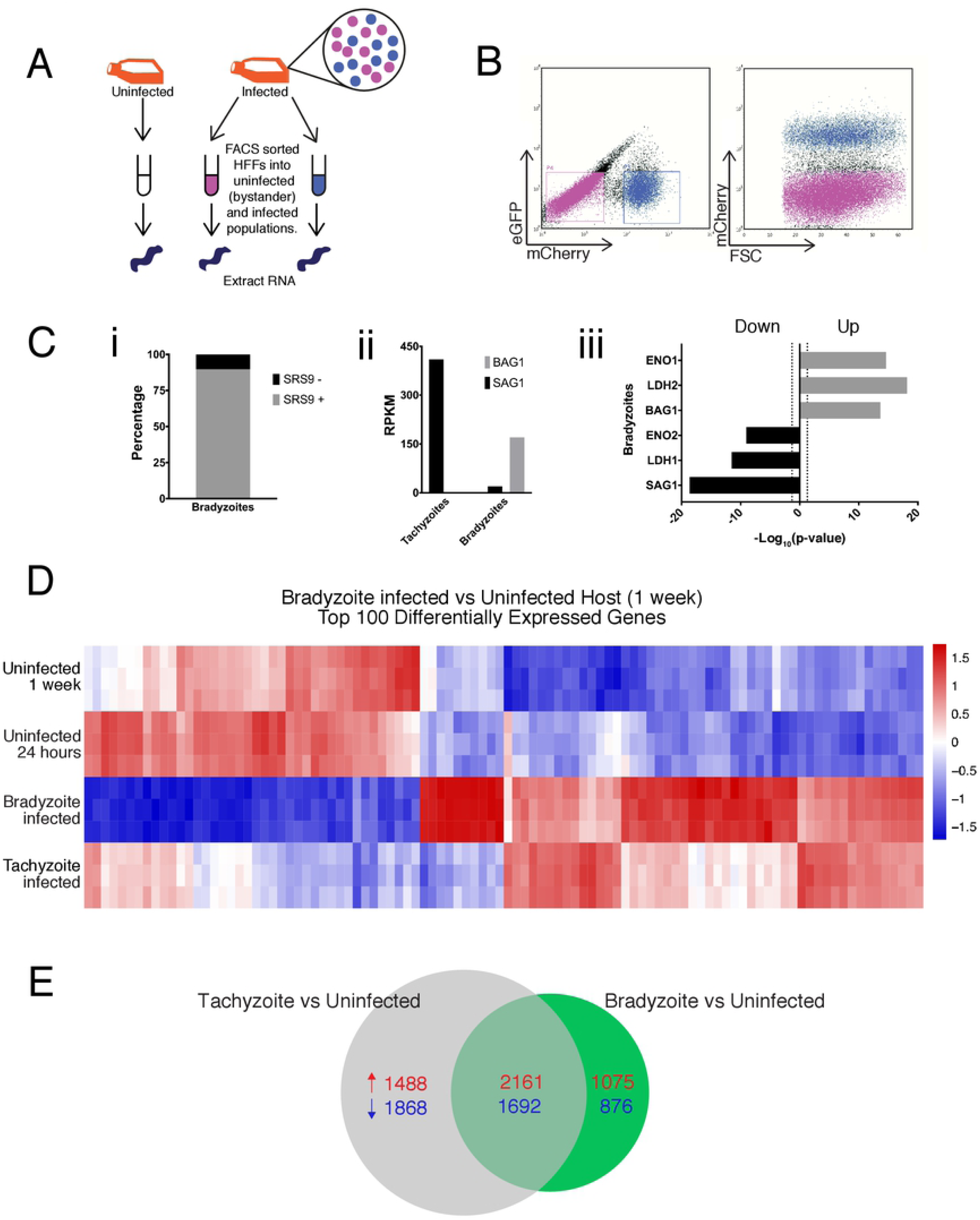
Transcriptional landscape of bradyzoite-containing host cells is different to tachyzoite infection. (**A**) Schematic of sample collection for RNA sequencing indicating uninfected, infected, and bystander populations. (**B**, *left*) a representative FACS plot of the sorting gates used to sort infected and bystander host cells defined as mCherry^+^eGFP^-^ (blue) and mCherry^-^eGFP^-^ (magenta) respectively; discarded cells indicated in black. (**B**, *right*) a FACS plot showing a shift towards a mCherry^+^ population after inoculation (**C**) (i) The proportion of SRS9^+^ (grey) and SRS9^-^ (black) parasites in bradyzoite samples. (ii) The levels of SAG1 and BAG1 reads (RPKM) in tachyzoite and bradyzoite samples. (iii) The negative Log_10_ of the P-value for known bradyzoite (grey) and tachyzoite (black) protein markers. (**D**) Inoculated HFF cells (human foreskin fibroblast) were FACS sorted into infected and uninfected samples before RNA extraction and sequencing (as per figure 1A) (n = 3 experiments, each with 3 replicates). Heat map of expression values for the top 100 differentially regulated genes between bradyzoite-infected (7 days *in vitro*) vs uninfected host cells (7 days). Top 100 DEGs expression are also shown for the tachyzoite-infected (24 hours) and uninfected host cells (24 hours) samples. Expression values are Log_2_ reads per kilobase per million (RPKM), scaled to have mean 0 and standard deviation 1 for each gene. The plots show that bradyzoites change the transcriptional profile of the host cell, and this appears to be somewhat different to the profile of tachyzoite infected host cells. See Table S1 for gene names. (**E**) Venn diagram of tachyzoite vs uninfected DEGs (24 hours post infection; grey) and bradyzoite vs uninfected DEGs (7 days post infection; green); the number of up-regulated genes are identified in red and down-regulated genes in blue.

Confident with the quality of our samples we then analysed the global effect of bradyzoites on the host transcriptome. This was done by comparing parasite-infected host cells with host cells from uninfected flasks. When bradyzoite-containing cells were compared with the uninfected HFF (after 1 week) we identified that 5804 host genes had significantly changed out of 13,793 expressed genes detected (∼42%), suggesting the presences of bradyzoites alters host cell transcription. This is slightly less than the number of differentially expressed genes upon tachyzoite infection (7209, ∼52%). To more quantitatively assess these changes we generated a heat map of the top 100 differentially expressed genes (DEG) for bradyzoite-infected cells verses uninfected host (1 week) and compared these to transcriptional profiles of the tachyzoite samples (Fig. 1D, see Table S1). From this set alone, similarities and differences in bradyzoite-containing and tachyzoite-containing host cells were evident. Some bradyzoite-induced changes were similar to those observed in tachyzoite infected host cells, whilst others appeared to be less pronounced or opposing, suggesting distinct transcriptional patterns between these two developmentally different stages (Fig. 1D, see Table S1, and Fig. 1E).

To further understand how host cells containing bradyzoites differ from tachyzoite infected cells, we wanted to determine how the changes in gene expression impacted host biological pathways. We first analysed all DEG of both bradyzoite- and tachyzoite-containing cells via over-representation analyses on the Molecular Signatures Database (MSigDB) Hallmark gene sets. (see Fig. S1A). Overall we could see a high perturbation across all biological pathways, suggesting re-wiring of the host transcriptome in both stages. However, some of these transcriptional changes may be secondary to parasite presence e.g. responses to endogenously released cytokines or non-specific disturbances. In order to tease out biological processes related to the presences of bradyzoites, we separated DEG into those that are specific to either developmental stage and those that are shared (Fig. 2A and see Fig. S1B respectively). Here we found that 3853 DEGs are shared between tachyzoite infected cells and bradyzoite infected cells (41%; 2161 up-reg., 1692 down-reg.), leaving 3356 tachyzoite specific DEGs (1488 up-reg., 1868 down-reg.), and 1951 DEGs specific to bradyzoite infection (1075 up-reg., 876 down-reg.) (Fig. 1E). We then performed over-representation analyses on each group. The majority of pathways in the Hallmark series showed a significant enrichment in the number of genes altered by both bradyzoites and tachyzoites (see Fig. S1B and Table S2). However, when the analyses for bradyzoite and tachyzoite specific DEG were compared, certain pathways are more enriched beyond the shared pathways (Fig. 2A, see Table S2). For an example, the *E2F*- and *Myc-targets* pathways were altered by tachyzoite infection, but no significant changes were seen in bradyzoite-containing cells over those that are shared (Fig. 2A and 2B, see Table S3). On the other hand, cells containing bradyzoites show a greater enrichment of DEGs in immune pathways, including the *IFN alpha-* and *IFN gamma-* response pathways, over what is already shared with tachyzoite infected cells (Fig. 2A and C, see Table S3). Overall, this implies that the presences of bradyzoites results in modulations of the host cell transcriptome, which is unique when compared to tachyzoites.

**Figure 2.**
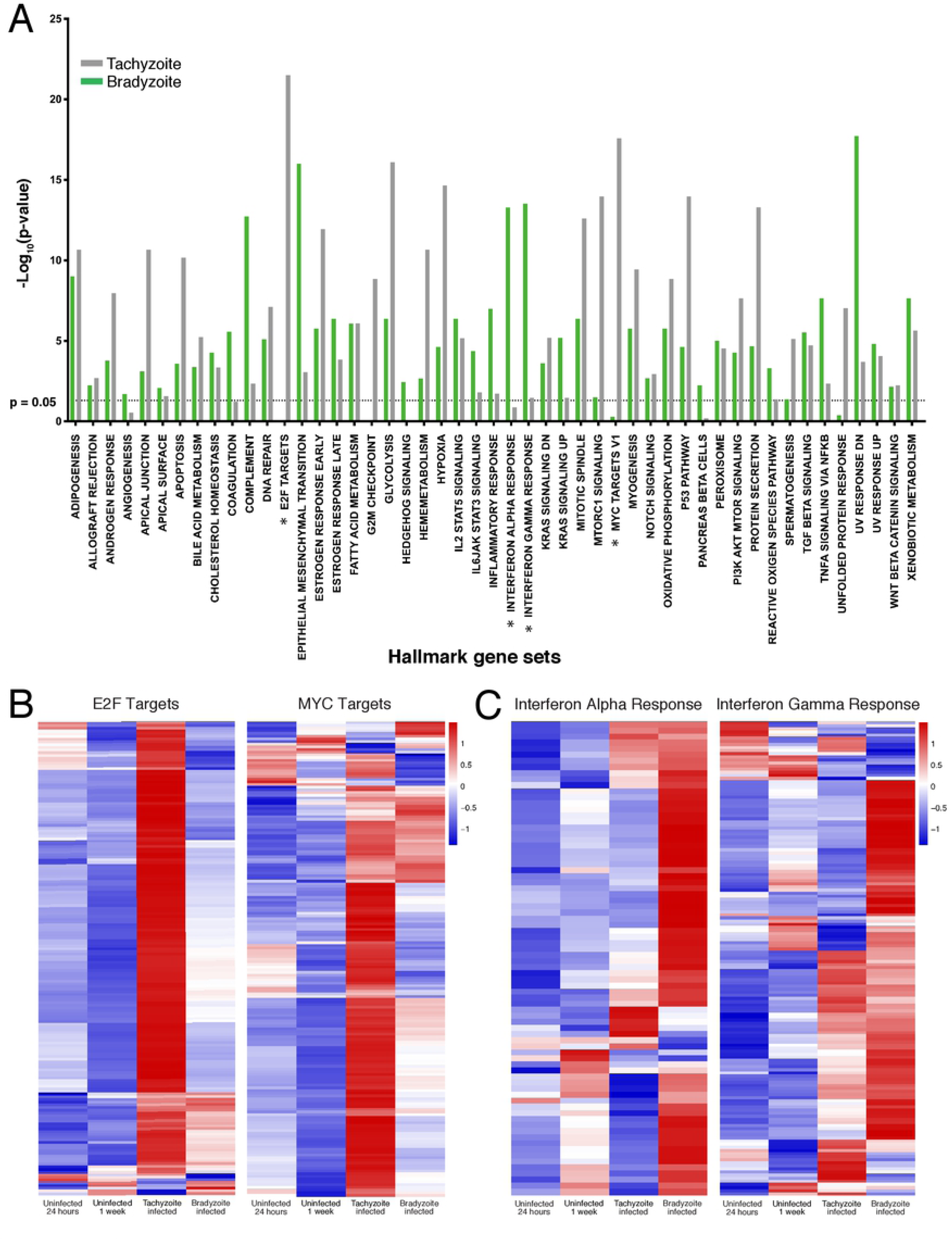
Tachyzoites and Bradyzoites modulate different host pathways. (**A**) Tachyzoites (grey) and bradyzoite (green) over-representation analysis results for each Hallmark gene set displayed as the negative Log_10_ of the P-value, dotted line p = 0.05. Stars indicate pathways with largely different profiles between tachyzoite and bradyzoites, which are shown in Figure 2B and C. (**B**) Pathways enriched during tachyzoite infection. Heat map of expression values for the genes that make up the *E2F target* and *MYC target* pathways. Expression values are Log_2_ RPKM, scaled to have mean 0 and standard deviation 1 for each gene. The plots show that tachyzoites alter the transcriptional profile of pathways *E2F target* and *MYC target* more so than bradyzoites. (**C**) Pathways enriched during bradyzoites infection. Heat map of expression values for the genes that make up the *Interferon Alpha Response* and *Interferon Gamma Response* pathways. Expression values are Log_2_ RPKM, scaled to have mean 0 and standard deviation 1 for each gene. The plots show that bradyzoite infections alters the transcriptional profile of pathways these two pathways more than bradyzoites. See also Figure S1, and Table S2 and Table S3 for gene lists.

### MYR1-dependent protein export plays a key role in host transcription modification in bradyzoites-containing host cells

Transcriptional changes can be imparted directly by pathogen effectors or as a result of downstream signalling after immune activation. We wished to specifically identify transcriptional changes induced by bradyzoites and to determine the overall effect of protein export on the host. To do this, we first determined whether MYR proteins, putative components of the PVM translocon and required for effector export, are present in bradyzoites^25, 28^. We assessed the relative expression of *myr1*, *2* and *3* in bradyzoites and tachyzoites from our RNAseq data (Fig. 3A). We found that *myr1, 2 and 3* are expressed in bradyzoites albeit at reduced levels (Fig. 3A). To functionally assess this complex in bradyzoites, we genetically tagged MYR1 with HA and could show that this protein is easily detectable and localises around the periphery of bradyzoites, similar to tachyzoites (see Fig. S2A), and is within the confines of the cyst wall, as marked by CST1 (Fig. 3B). We then knocked-out MYR1 in mCherry-expressing-Pru using CRISPR/Cas9 by targeting the first exon to ensure disruption. This resulted in a random 69bp insertion leading to three stop codons (see Fig. S2B) and was functionally confirmed by the inability of parasites to export transiently expressed GRA24-Myc (see Fig. S2C)^27^. Next, we assessed differentiation capacity and morphology of Δ*myr1* bradyzoites by inducing bradyzoite formation *in vitro*. There was no obvious differences in morphology indicated by an intact cyst wall, compact parasites expressing SRS9, and the presence of amylopectin granules (Fig. 3C (i)). The number of cyst-containing host cells was significantly less in Δ*myr1* samples (Fig. 3C (ii)) (see below for potential explanation of this); however, the size of the cysts was not altered by the absence of MYR1 (Fig. 3C (iii)). Overall, this suggests that cyst development is not critically reliant on MYR1, at least in tissue culture. This is in contrast to deletion of ASP5, which results in defects in cyst wall biosynthesis^30^.

**Figure 3.**
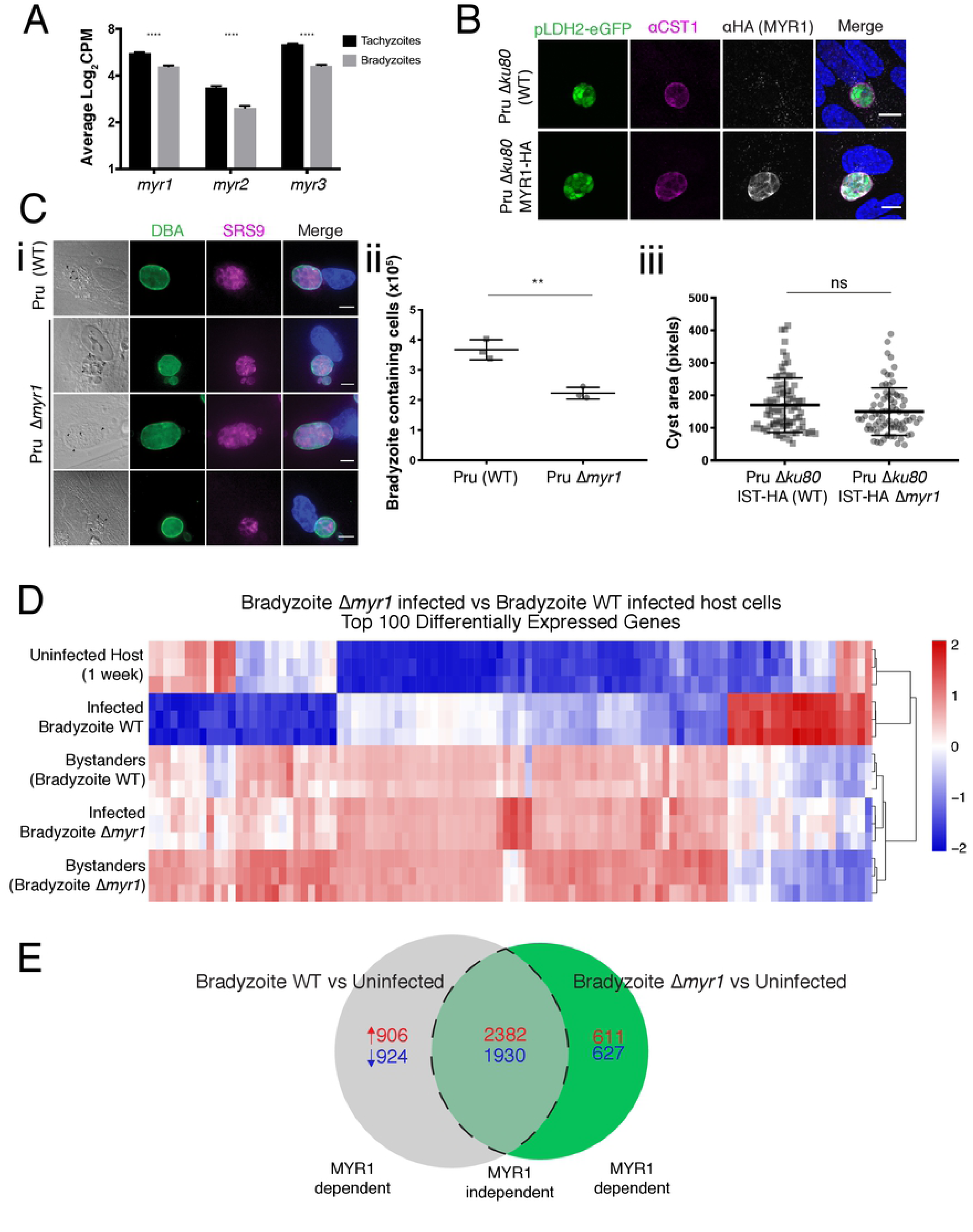
MYR1 plays a role regulating host cell transcriptional changes in bradyzoites containing cells. (**A**) Differences in the expressional profile (Log_2_CPM) of MYR1, MYR2, and MYR3 between tachyzoites (24 hours) and bradyzoites (7 days *in vitro*). Expression drops during bradyzoite stages for all proteins. (**B**) PruΔ*ku80* and PruΔ*ku80* MYR1-HA parasites were induced under alkaline conditions for 7 days before fixation. MYR1-HA (αHA; white) in bradyzoites (pLDH2-eGFP; green) is localised to the PV space and co-localises to cyst wall marker CST1 (αCST1; magenta). Brightness and contrast was edited on single colour channels prior to merging. (**C**) (i) WT and Δ*myr1* parasites were fixed after 7 days of differentiation under alkaline conditions. Both WT and Δ*myr1* were able to form intact cysts (DBA; green) containing health bradyzoites (αSRS9; magenta) and showed the presence of amylopectin granules (black arrows). Scale bar = 10μM; MYR, Myc Regulationr; PV, parasitophorous vacuole; DAPI, 4’,6-diamidino-2-phenylindole; DBA, fluorescein conjugated Dolichos biflorus agglutinin. Brightness and contrast was edited on single colour channels prior to merging. (ii) The number of bradyzoite containing cells after 7 days of differentiation between WT (Pru) and Δ*myr1* was analysed by FACS (mean ± SD, n = 3 experiments, unpaired, two-sided Student’s t-test with Welch’s correction; **p=0.0058). (iii). The size of PruΔ*ku80* Δ*myr1* cysts were compared to WT by quantification of from immuno-fluorescence images (mean ± SD, n = 3 experiments, each experiment analysed on average between 20-40 bradyzoite cysts, unpaired, two-sided Student’s t-test with Mann-Whitney correction; p=0.156) (**D**) WT (Pru) and Δ*myr1* inoculated human foreskin fibroblast (HFF) cells were FACS sorted into infected and uninfected samples before RNA extraction and sequencing (as per Fig. 1A). Heat map of log_2_RPKM expression for the top 100 differentially regulated genes between WT bradyzoite infected and Δ*myr1* bradyzoite infected cells. Expression has been scaled to have mean 0 and standard deviation 1 for each gene. In the absence of MYR1, bradyzoite specific host expression appears to somewhat mimic the bystander cell population (from both WT and Δ*myr1*). See also Figure S2 and Table S4 for gene list. (**E**) Venn diagram of bradyzoite WT infected vs uninfected DEGs (grey) and bradyzoite Δ*myr1* infected vs uninfected DEGs (green); the number of up-regulated genes are identified in red and down-regulated genes in blue. Dotted lines indicated shared DEGs deemed as independent of MYR1.

The ability to form healthy Δ*myr1* cysts then enabled us to determine what transcriptional changes are specifically caused by potential bradyzoite effectors in the early stages of chronic infection (7 days), compared to indirect effects such as immune signalling. Following the procedure laid out in Figure 1, uninfected, bradyzoite-containing and uninfected bystander cells were collected. When WT-infected cells were compared with Δ*myr1*-infected cells, we observed dramatic differences in gene expression in the absence of MYR1. A heat map generated of the top 100 DEG identified between WT-infected and Δ*myr1*-infected cells shows that the loss of MYR1 results in a transcriptional profile that closely mirrors the transcriptional changes seen in bystander cells (Fig. 3D, see Table S4). Reinforcing this concept, Δ*myr1-*infected cells were more closely related in the hierarchical heatmap with bystander cells rather than WT-infected cells (whereas bradyzoite-WT-containing and uninfected cells are excluded from this cluster). Moreover, bystander cells are vastly different to uninfected cells, implying that uninfected bystander cells are impacted by the presence of bradyzoites in culture. Together these data suggests that transcriptional changes observed in the presence of bradyzoites are actively modulated and that protein export via MYR1 plays a role in altering the transcriptional program during chronic infection.

We then assessed what impact MYR1-dependent protein export plays in altering host biological pathways of bradyzoite-containing cells. We first compared the DEG analysis of WT-infected vs uninfected with Δ*myr1*-infected vs uninfected and found the expression of 3068 host genes (∼42%) was dependent on MYR1 (Fig. 3E). The remaining genes (4312) were altered in the same direction in both comparisons; however, a limitation with this type of analysis is that it does not distinguish between the degree of change. For an example, XIAP is significantly upregulated in both WT infected cells and Δ*myr1* infected cells when compared to uninfected host, but to different degrees which is seemingly dependent on MYR1 (see Fig. S3C). Nevertheless, it is evident that MYR1 has an impact on the host transcriptional response to the presence of bradyzoites.

We performed over-representation analyses using the Hallmark gene sets on each group and observed that bradyzoites depend on MYR1 in order to alter these pathways. In accordance with the first data set, it is evident that the presences of bradyzoites results in a multitude of changes in the host transcriptome, which impact all the Hallmark pathways. All the pathways examined contain a significant number of genes that change in expression following the loss of MYR1; however, the modulation of some pathways is less dependent on MYR1 than others. For example, pathways *TNF-alpha signalling via NFKB* and *Epithelial Mesenchymal Transition* are highly enriched in the MYR1 independent gene set (see Fig. S3B and Table S5), whereas the manipulation of other pathways such as *Adipogenesis* and *Glycolysis* are more dependent on MYR1 (see Fig. S3A and Table S5). Nevertheless, differentially expressed genes dependent on MYR1 are present in the majority of the pathways to a significant degree.

These data represents all the gene expression changes upon bradyzoite infection, including those that are present during tachyzoite infection. To determine what role protein export plays specifically in bradyzoite host manipulation, we examined the Hallmark pathways that saw a significant enrichment of DEGs specific to bradyzoites (Fig. 1E) and determined the proportion of the pathway that was dependent on MYR1. We found that all pathways were significantly altered by the absence of MYR1 (Fig. 4, see Table S5). This shows that a large proportion of bradyzoite induced host transcriptome manipulation requires the presence of MYR1, and further implicates the importance of active export of effector proteins in bradyzoites.

**Figure 4.**
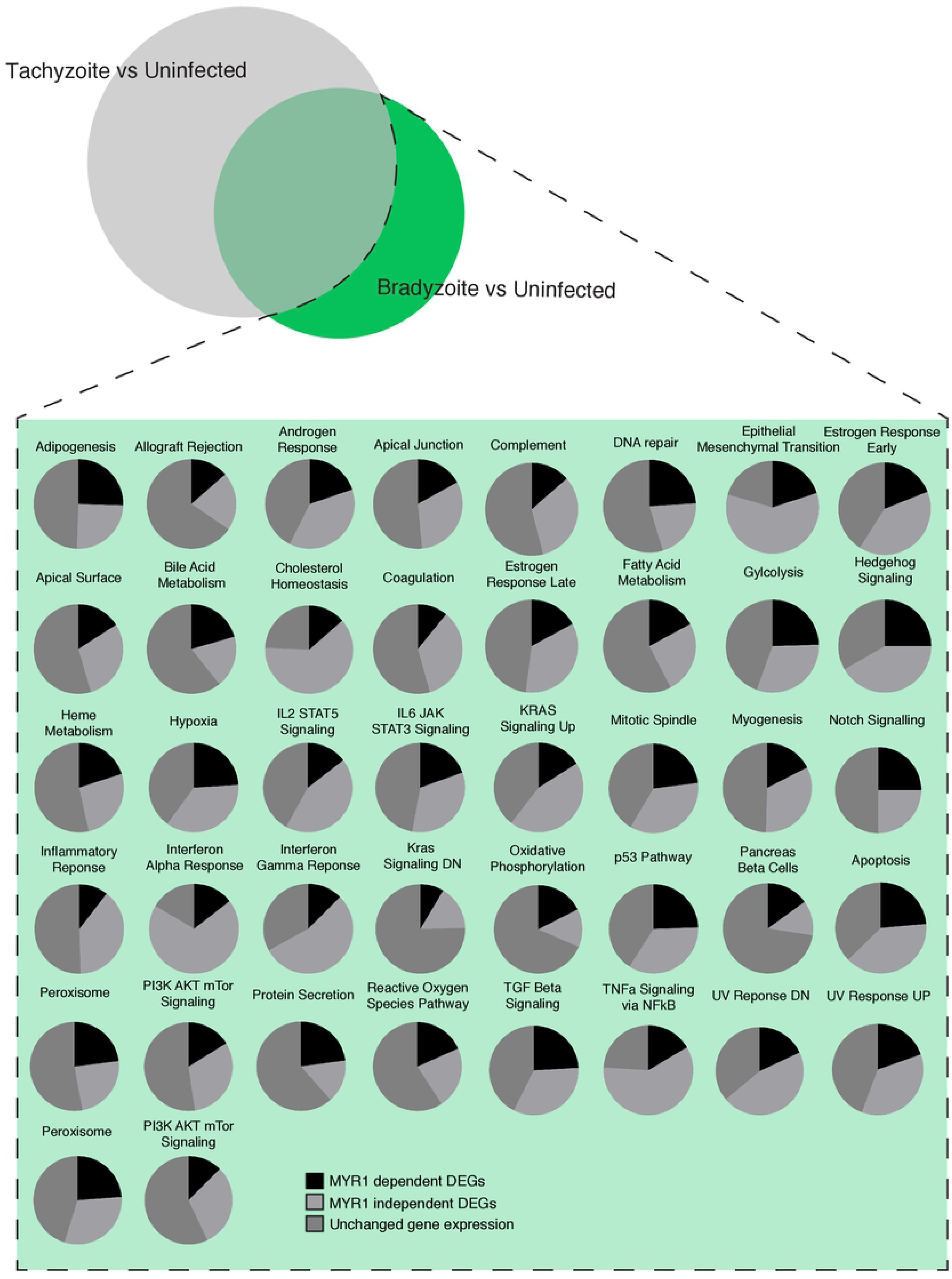
Proportion of bradyzoite specific pathways changes are dependent on MYR1. Bradyzoite specific DEGs shown in Fig. 2 was used for GSEA Hallmark gene set pathway analysis. The proportion of DEGs that were dependent on MYR1 (as determined by Fig. 4B) were then shown in those pathways that were significantly altered in bradyzoites (from Fig. 2B). Black, MYR1 dependent DEGs; Light grey, MYR1 independent DEGs; Dark grey, unchanged DEGs. See also Figure S3 and Table S5 for gene list.

### IST is expressed and exported in bradyzoites

Given our findings above, we hypothesised that bradyzoites have a repertoire of effector proteins that are exported into the host, some of which may also be exported in tachyzoites. After several unsuccessful attempts at determining, in an unbiased fashion, bradyzoite specific effector proteins, we focused on known effector proteins, GRA16, GRA24 and IST^17–20^. We first compared the proportion of reads between the infected bradyzoite and tachyzoites samples, from the data in our original RNAseq analysis. We saw a significant drop in the expression of GRA16 (Log_2_ fold change - 2.681) and GRA24 (Log_2_ fold change −2.159) in bradyzoite samples, whereas IST appears to be slightly more expressed (Log_2_ fold change +0.374) (Fig. 5A). To validate that mRNA expression changes were translatable to protein expression, we HA tagged these proteins in PruΔ*ku80* and probed for their expression via western blot. GRA16, GRA24 and IST were all expressed in tachyzoites, whereas bradyzoites (classified by SRS9 expression and SAG1 absence) expressed GRA16 and IST (Fig. 5B). This indicated that at least two known effectors are present in bradyzoites.

**Figure 5.**
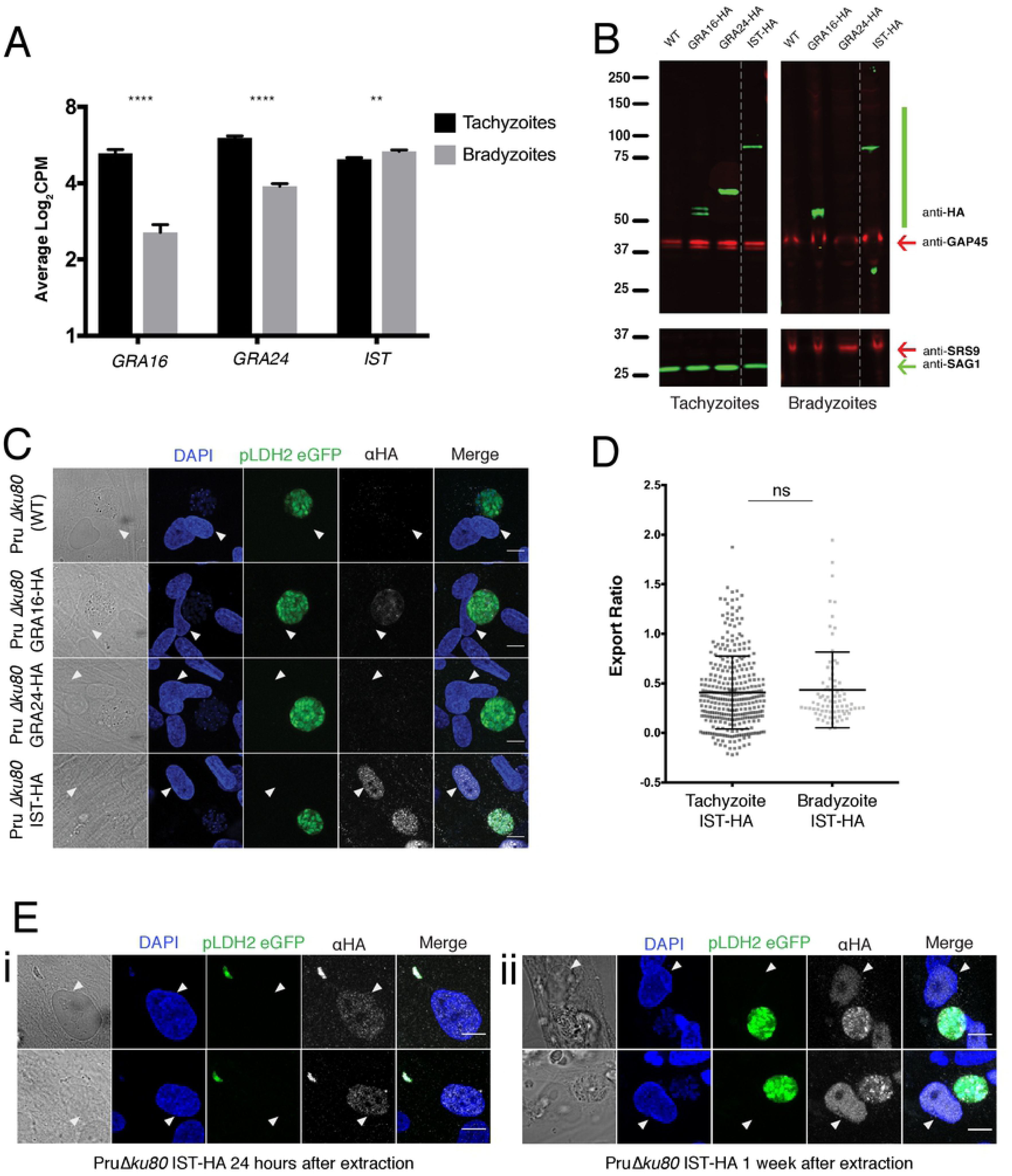
Bradyzoites produce and export IST. (**A**) Differences in the expressional profile (Log_2_CPM) of *GRA16*, *GRA24*, and *IST* between tachyzoites (24 hours) and bradyzoites (7 days *in vitro*). *GRA16* and *GRA24* reads drop during bradyzoite stages, whereas *IST* levels slightly increase (n = 3 experiments, each with 3 replicates; ****p<0.0001) (**B**) Western blot indicates GRA16-HA, GRA24-HA and IST-HA (PruΔ*ku80* background) are expressed in tachyzoites (lysed), whereas only GRA16-HA and IST-HA are expressed in bradyzoites (7 days in vitro). αGAP45 is used as a loading control, αSAG1 and αSRS9 are used as tachyzoite and bradyzoite controls respectively. (**C**) PruΔ*ku80* WT, GRA16-HA, GRA24-HA and IST-HA parasites were differentiated under alkaline conditions for 7 days before fixation. GRA24-HA (αHA; white) was not produced by bradyzoites (pLDH2-eGFP; green) or exported into the host nucleus. In contrast, bradyzoites were able to produce GRA16-HA and IST-HA; however, only IST-HA was exported into the host nucleus. White arrows indicate location of host nucleus (DAPI; blue). (**D**) No change in the ratio (nuclear intensity/vacuolar intensity) of IST export between tachyzoites and bradyzoites from immuno-fluorescence imaging (Figure 6c) (mean ± SD, n = 3 experiments, each experiment analysed on average between 20-40 bradyzoite cysts and 60-200 tachyzoite vacuoles; unpaired, two-sided Student’s t-test; ****p<0.0001). (**E**) PruΔ*ku80*IST-HA parasites were differentiated under alkaline conditions for 7 days before extracted and reinoculated. Samples were either fixed 24hours (i) or 1 week post infection (ii). IST is consistently exported during bradyzoites infection. Scale bar = 10μM. Brightness and contrast was edited on single colour channels prior to merging. Scale bar = 10μM; GRA, dense granule; IST, inhibitor of STAT1 transcriptional activity; PV, parasitophorous vacuole; DAPI, 4’,6-diamidino-2-phenylindole. Brightness and contrast was edited on single colour channels prior to merging. See also Figure S4.

In order to establish if GRA16 and IST are found in the host nucleus in bradyzoite-containing cells, we quantitated their expression and localisation by microscopy (Fig. 5C). GRA24 is consistently not expressed in 7-day-old bradyzoites; accordingly, the presence of GRA24 in the host nucleus is comparable to un-tagged WT controls (Fig. 5C and S4A). Interestingly, there appears to be some GRA16 present within the bradyzoite PV, but it is undetectable in the host nucleus (Fig. 5C and S4A). In contrast, we observed the export of IST into the host nucleus by bradyzoites (Fig. 5C and S4A) in equivalent amounts to tachyzoites (Fig. 5D). Therefore the increase of expression does not directly indicate an increase in the amount of protein exported into the host cell. To resolve whether the presence of IST in the host nucleus is residual protein from tachyzoite stages or is synthesised and exported in bradyzoites, we differentiated bradyzoites for a week, before extracting via needle passage and inoculating new host cells. Both 24 hours and one week after inoculation, we continue to see IST export in bradyzoites (Fig. 5E(i) and (ii)). This strongly suggests that IST is produced in and exported by bradyzoites.

Following the identification of IST as a bradyzoite exported effector protein, we sought to determine whether MYR1 was also responsible for IST export in bradyzoite stages and whether this effector was functional in controlling IFNγ-dependent responses. To investigate this, we created knockouts of both *myr1* and *ist* in the PruΔ*ku80* IST-HA line by coupling CRISPR/Cas9 with an homologous template containing the DHFR selectable marker followed by verification via western blot and microscopy, respectively (see Fig. S5A and B). Upon *myr1* disruption, the export of IST in bradyzoites is significantly inhibited (Fig. 6A(i) and (iiI)) and is comparable to background levels of cells infected with Δ*ist* and WT PruΔ*ku80* levels (Fig. 6A(i) and (ii)). Together, these data show that bradyzoites differentially express and actively export known effector protein, IST, via MYR1 during infection.

**Figure 6.**
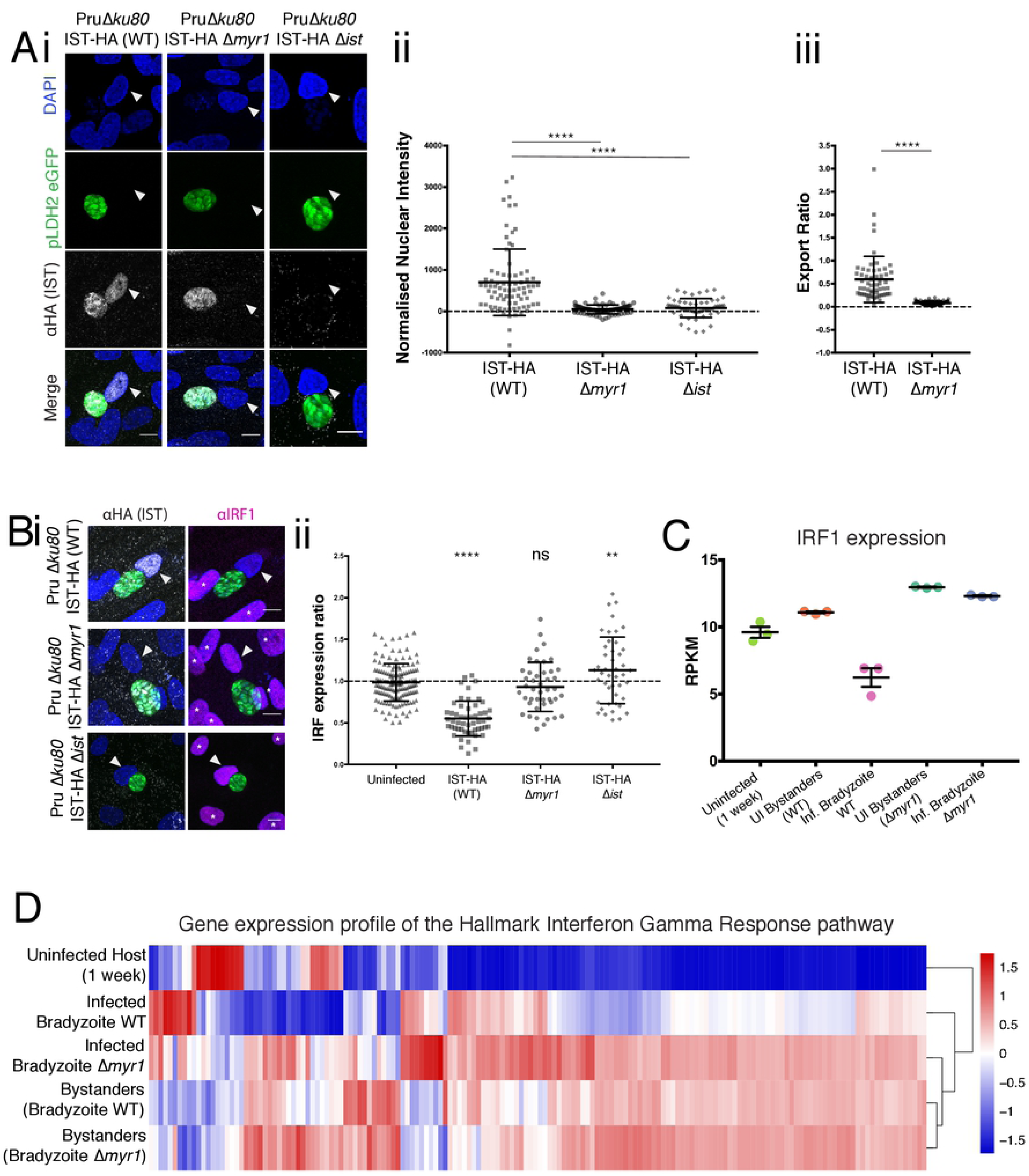
Bradyzoites alter the Interferon gamma pathway. (**A**) (i) PruΔ*ku80* IST-HA (WT), PruΔ*ku80* IST-HA Δ*myr1*, and PruΔ*ku80* IST-HA Δ*ist* parasites were differentiated under alkaline conditions for 7 days before fixation. The absence of MYR1 in bradyzoites (pLDH2-eGFP; green) inhibits the export of IST-HA (αHA; white) into the host nucleus. White arrows indicate location of infected host nucleus (DAPI; blue). White stars indicate location of un-infected host nuclei. Scale bar = 10μM. Brightness and contrast was edited on single colour channels prior to merging. (ii) The increase in IST-HA mean fluorescence intensity in the host nuclei of WT infected cells is lost in Δ*myr1* and Δ*ist* infected host cells (normalised to PruΔ*ku80* untagged controls). Analysed from immuno-fluorescence imaging (Figure 7ai) (mean ± SD, n = 3 experiments, each experiment analysed on average between 20-30 bradyzoite cysts; unpaired, parametric one-way ANOVA test with Dunn’s multiple comparison; ****p<0.0001). (iii) The ratio (nuclear intensity/vacuolar intensity) of IST drops in Δ*myr1* when compared to WT. Analysed from immuno-fluorescence imaging (Figure 7ai) (mean ± SD, n = 3 experiments, each experiment analysed on average between 20-30 bradyzoite cysts; unpaired, two-sided Student’s t-test; ****p<0.0001). (**B**) (i) PruΔ*ku80* IST-HA (WT), Δ*myr1*, and Δ*ist* parasites were differentiated under alkaline conditions for 7 days before being treated with 50ng/ml of human-IFNγ for 15hrs and fixed. Nuclear IRF1 (αIRF1; magenta) protein levels drop to varying degrees in WT infected cells, and this inhibition is lost in Δ*myr1* and Δ*ist* bradyzoite (pLDH2-eGFP; green) infected host cells. White arrows indicate location of host nucleus (DAPI; blue). White stars indicate location of un-infected host nuclei. Scale bar = 10μM. Brightness and contrast was edited on single colour channels prior to merging. (ii) The inhibition of IRF1 by WT bradyzoites is reversed by the knockout of *myr1* and IRF1 levels return to similar levels when compared to uninfected and Δ*ist* infected cells. Analysed from immuno-fluorescence imaging and normalised to surrounding uninfected cells (Figure 7bi) (mean ± SD, n = 3 experiments, each experiment analysed on average between 20-30 bradyzoite infected host cells; unpaired, parametric one-way ANOVA test with Dunn’s multiple comparison; **p<0.01, ****p<0.0001). (**C**) RPKM values for IRF1 from Figure 3 data set. (**D**) A gene heat map of log_2_RPKM expression for the Hallmark gene set *IFN gamma response* showing all samples from the Figure 3 analysis. Expression has been scaled to have mean 0 and standard deviation 1 for each gene. See also Figure S5, S6 and Table S6 for gene list.

### Bradyzoites supress the Interferon gamma pathway through MYR1-dependent IST export

In tachyzoite infection, IST has been shown to inhibit the host cell response to IFNγ stimulation^19, 20^. To determine if this holds true for bradyzoite infection, we stimulated bradyzoite-containing cells with IFNγ and probed for IRF1, a downstream protein which is upregulated upon IFNγ stimulation and pSTAT1-GAS binding^76^. As is the case in tachyzoites, IRF1 expression is significantly inhibited in cells containing WT cysts as compared to bystander cells (Fig. 6B(i) and (ii)). On the other hand Δ*myr1* and Δ*ist* parasites are unable to inhibit IRF1 expression and are comparable to uninfected bystander cells (Fig. 6B(i) and (ii)). The significant lower levels of IRF1 expression in Δ*myr1* as compared to Δ*ist* could be explained by an underestimation due to host cell death, as discussed below. Interestingly, we noticed that the ability for a cyst to inhibit IRF1 expression varied widely between host cells, from having no effect to complete inhibition (Fig. 6B(ii) and S5C(i); range 1.069-0.133). We checked whether this variation could be explained by the correlation between the amount of IST in the nucleus and the subsequent expression of IRF1. Our observations could be explained by fitting a hyperbolic curve (blue; see Fig. S5C(ii)) (*p* = 1.042×10^−12^). As the hyperbolic curve indicates, the level of IRF1 decreases as amount of IST-HA in the nucleus increases; however, there is a point at which high levels of IST cannot further decrease IRF1 expression. The variability in parasite induced IRF1 inhibition was also noted in the tachyzoite stages of the PruΔ*ku80* strain, with no significant difference in the degree of inhibition between the two stages (see Fig. S5D). Interestingly, RHΔ*ku80* strains showed a stronger ability to suppress IRF1 with less variability (see Fig. S5D).

To further analyse the role of MYR1 (and therefore indirectly IST) in the host cell’s response to bradyzoite infection, we returned to our dataset derived from Δ*myr1* bradyzoites (Fig. 3 and 4). Despite the absence of IFNγ stimulation in our RNA sequencing data, we observed a difference in the expression of IRF1 between WT infected and Δ*myr1* infected cells. In the absence of *myr1*, the transcript levels of IRF1 is comparable to that of uninfected bystander cells, as previously shown by Olias et al., (2016) (Fig. 6C). This pattern is consistent in several genes previously identified via CHIP-Seq analysis as IFNγ-induced pSTAT1-GAS transcripts, including GBPs (see Fig. S6)^77^. A gene expression heat map of the Hallmark gene set: *Interferon gamma response* (Fig. 6D, see Table S6) illustrates that Δ*myr1* infected cells cluster more closely with uninfected bystander cells than with WT infected cells, suggesting that the majority of transcriptional changes that occur within the *Interferon gamma response* pathway are dependent on MYR1 in bradyzoites. Overall, this work indicates that bradyzoites are altering the host cells response to IFNγ stimulation via the export of IST through the MYR1 pathway.

### Protein export is important for survival of bradyzoite-containing cells upon IFNγ challenge

During our experiments, we noticed that the number of intact Δ*myr1* cysts was greatly reduced after IFNγ treatment, suggesting this cytokine may cause death of infected host cells. To understand this in detail, we performed live imaging of WT, Δ*myr1* and Δ*ist* over 15 hours following IFNγ treatment (Fig. 7A (i); see examples movies S1 (WT), S2 (Δ*myr1*) and S3(Δ*ist*)) and monitored host cell death by Propidium Iodide (PI) uptake. Upon IFNγ treatment we observed that there was a significant decrease in the survival of infected host cells when compared to negative controls (Fig. 7A(ii)). Cells containing Δ*myr1* cysts had the largest decrease in cell survival (87%), whereas WT and Δ*ist* saw a drop of only 28% and 37%, respectively. This equated to no significant difference between IFNγ treated WT and Δ*ist* cyst numbers at the end of the 15 hour period, whilst loss of MYR1 was significantly different from both WT and Δ*ist*, suggesting a role for other unknown MYR1-dependent exported protein/s in blocking IFNγ-induced host cell death (Fig. 7A(ii)). Analysing the same samples using FACS and monitoring GFP expressed from the bradyzoite-specific LDH2 promoter also showed a similar outcome (Fig. 7B(i) & (ii))^75^. In stark contrast, IFNγ treatment of cells infected with Δ*myr1-*tachyzoites did not have a significant cell death phenotype, strongly supporting the notion that our bradyzoite-infected cells are functionally different from tachyzoite-infected cells (see Fig S7A). These data suggest that IST alone is not accountable for the large volume of cell loss seen by Δ*myr1*-bradyzoite-containing cells, implying other factors contribute to prevent cell death.

**Figure 7.**
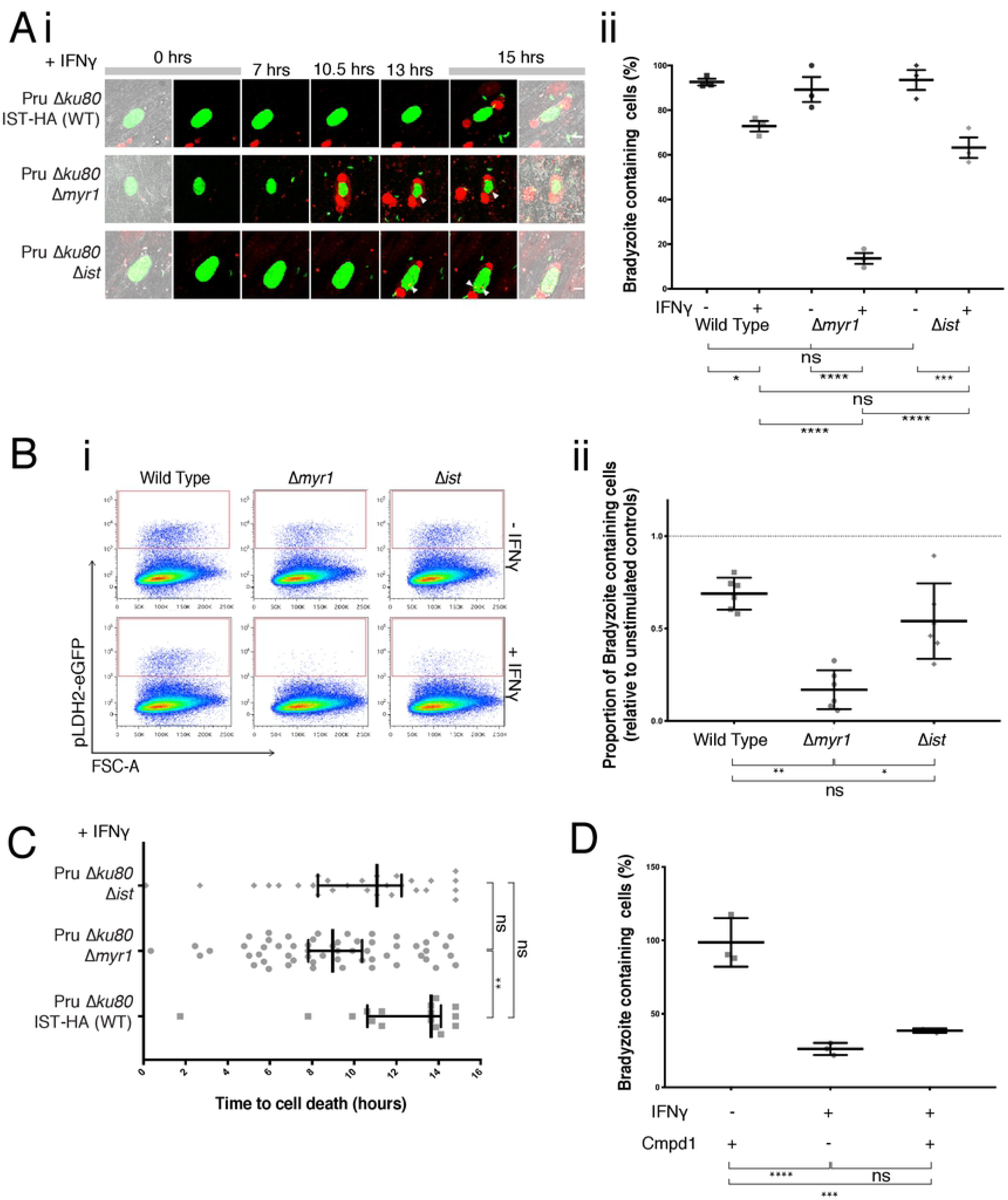
Depletion of MYR1 prevents cyst persistence and induces host cell death upon IFNγ stimulation. (**A**) (i) PruΔ*ku80* IST-HA (WT), PruΔ*ku80* IST-HA Δ*myr1*, and PruΔ*ku80* IST-HA Δ*ist* parasites were differentiated under alkaline conditions for 7 days before imaging. During each experiment (n=3), between 15-35 parasite cysts were imaged every 14 mins for 15 hours after the addition of PI (red) and 50ng/ml of IFNγ. PI can be seen entering the host cell and bradyzoites (pLDH2-eGFP; green; white arrow) are released. Scale bar = 10μM. Brightness and contrast was edited on single colour channels prior to merging. (ii) The percentage of cyst infected cells that survived the 15 hours treatment of IFNγ drops by almost 90% in Δ*myr1* infected cells, significantly more than both WT and Δ*ist* (mean ± SD, n = 3 experiments, each experiment tracked between 15-30 bradyzoite cysts; unpaired, one-way ANOVA test with Sidak’s multiple comparison test; *p<0.05, ***p<0.001, ****p<0.0001). (**B**) (i) PruΔ*ku80* IST-HA (WT), Δ*myr1*, and Δ*ist* parasites were differentiated under alkaline conditions for 7 days before being treated with 50ng/ml of human-IFNγ for 15hrs, trypinised and FACS sorted. FACS plot showing gating of eGFP^+^ cells (red) of WT, Δ*myr1*, and Δ*ist* bradyzoite infected host cells. (ii) Proportion of bradyzoite-eGFP^+^ cells compared to unstimulated samples shows a similar pattern to the live imaging data from Figure 8a. (mean ± SD, n = 6 experiments, unpaired, one-way ANOVA test with Sidak’s multiple comparison test; *p<0.05, **p<0.01). (**C**) Time to cell death in hours; Δ*myr1* infected cells died earlier than both WT and Δ*ist* (median ± 95% CI, n = 3 experiments, each experiment tracked between 15-30 bradyzoite cysts; unpaired, nonparametric one-way Kruskal-Wallis ANOVA test with Dunn’s multiple comparison; **p<0.01). (**D**) Compound 1 (2μM) was added to 7-day-old bradyzoites for an hour prior to the IFNγ addition for 15 hours. Samples were fixed, FACS analysed, and the proportion of bradyzoite-eGFP^+^ cells compared to untreated samples (mean ± SD, n = 3 experiments, unpaired, one-way ANOVA test with Sidak’s multiple comparison test; ***p<0.001, ****p<0.0001, p=0.510).

We then observed the morphology of cysts over the filming period and tracked how long it took for cell death to occur. Here we could see that cell death was accompanied by PI uptake, which was observed as staining of the host nucleus and debris. During cell death, host cells contracted before cell blebbing occurred (Fig. 7A(i), and SV1, SV2, SV3 in the supplementary material), a typical sign of programmed cell death. For Δ*myr1* infected cells, cell death occurred earlier, at approximately 9 hours post IFNγ stimulation, and was significantly earlier than those WT-infected cells that died (13.65 hours) (Fig. 7C). Interestingly, the time of death for Δ*ist*-infected cells was between Δ*myr1*- and WT-infected cells (at approximately 11 hours), further implying that IST alone cannot account for the host cell response to IFNγ during infection. We also observed that although some bradyzoites are released from the cyst, a majority remain within the dying cell and in some cases parasites are also observed taking up PI (Fig. 7A(i); white arrows).

We wanted to determine if this phenomenon was host cell driven or if it was caused by the egress of parasites from host cells. The latter would suggest bradyzoites can sense a change in the environment upon IFNγ treatment resulting in egress activation, whereas the former implies the host cell initiates cell death in the absence of parasite modulation. To disentangle these two pathways we used compound 1, which has been shown to prevent egress by inhibiting PKG^61, 78^. We hypothesised that if parasite egress was responsible for the cell death, then the addition of compound 1 would result in the recovery of the phenotype. Bradyzoite samples were incubated with compound 1 for an hour prior to the addition of IFNγ and 15 hours later checked for the levels of bradyzoite-containing host cells. Compound 1 was able to inhibit bradyzoite egress (see Fig. S7C); however, there was no significant difference in cell survival between cells treated with and without compound 1 upon IFNγ treatment (Fig. 7D). Furthermore, PI uptake was observed in bradyzoite-containing cells treated with compound 1 and IFNγ suggesting cell death was still occurring in the absence of parasite lysis (see Fig. S7B; white arrows indicate PI-positive parasites). These data indicate that one or more MYR1-dependent effectors protect bradyzoite-containing host cells from undergoing IFNγ-mediated cell death, demonstrating the importance of host cell modification during immune challenge, at least in early bradyzoite forms.

## Discussion

Chronic *Toxoplasma* infection can result in progressive blindness, act as a reservoir for disease reactivation, is a source of transmission in the food chain^1^ and more recently has been associated with several psychiatric disorders^4^. The interactions between *Toxoplasma* and its host cell are likely critical to the parasites ability to persist for life and alter the physiology of the brain. Our study shows, for the first time, that bradyzoites actively manipulate their host cell, which is functionally relevant to persistence upon immune challenge. We have established that host cells containing 7 day-old bradyzoites have a transcriptional profile that is different from acute-stage tachyzoite forms. We show that MYR1 (a likely component of the export machinery) is expressed in bradyzoites and that its absence leads to an inability to induce transcriptional changes in bradyzoites containing cells. Furthermore, we show that IST, expressed from its native promoter, is exported by bradyzoites into host cells in a MYR1-dependent fashion, thus strongly suggesting that bradyzoites are capable of protein export into and modification of their host cell. Given that IST represses STAT1 signalling during IFNγ stimulation^20^, our data suggests that IST could be important for bradyzoite persistence in muscle and CNS tissue, a line of enquiry future *in vivo* studies could focus on.

Our data is somewhat at odds with a recent study by Krishnamurthy and Saeij (2018)^79^, which shows that the enforced expression of two tachyzoites effectors in bradyzoites (GRA16 and GRA24) does not result in export. Instead these effectors accumulated within the cyst wall, therefore suggesting that protein export might not occur at these stages. The authors suggest this could be because these proteins are not required during chronic stages, and either the cyst wall creates a physical barrier to protein export or that bradyzoites lack a translocation complex (i.e. MYR1). We, on the other hand, monitor all protein export from native promoters and we show here that GRA24 is not expressed, whilst GRA16 appears to be expressed but not exported. In the case of GRA16, the experiments reported here are in line with Krishnamurthy and Saeij’s study and the reason for production of this protein without export is yet underdetermined. Interestingly, in bradyzoites we see an increase in IST expression as well as a similar level of export, when compared to tachyzoites. IST was detected in host cells that were both infected with tachyzoites and subsequently differentiated into bradyzoites for 7 days, and in samples where isolated bradyzoites were re-infected onto new host cells. Together with our transcriptional data showing increased expression of this effector we are confident that bradyzoites are capable of exporting IST into the host cell nucleus. What our data does not address is whether translocation across the cyst wall occurs. It remains possible from our study that all IST observed in the host cell nucleus of 7 day old bradyzoites is exported before the production of the cyst wall. For this to be true IST would have to be extremely stable and have a very low turnover rate; this would enable IST to stay at the levels we observed in tachyzoites after 7 days of differentiation. The other possibility is that there is something unique about IST that allows for its export across the cyst wall in bradyzoites but not GRA16 and GRA24. The only known trafficking element in exported proteins is the TEXEL motif^27, 29^, but this cannot explain the export differences observed given this motif is present in both GRA16 and IST^17, 19^. If this was the case then IST should be able to be exported when expressed under the inducible promoter that Krishnamurthy and Saeij use^79^.

We further show that the transcription factor IRF1, which is expressed in a IFNγ dependent fashion, is blocked by the presence of IST. This suggests that, at least at the early timepoints (7 days), bradyzoites do have the ability to block IFNγ signalling just as well as tachyzoites. However, our data also indicates that IST alone does not appear to protect bradyzoite-containing host cells from IFNγ-mediated cell death.

The lifestyles of tachyzoites and bradyzoites are vastly different and thus they have distinct requirements of their host cell. Tachyzoites are fast growing and aim to spread through tissue and around the body. Bradyzoites, on the other hand, are slow growing and aim to persist for the life of their host, seemingly avoiding immune detection. Here we show that bradyzoite-containing host cells have a distinct transcriptional landscape, which differs from acute stage tachyzoites perhaps reflecting specific survival requirements. Tachyzoites appear to impact the host’s cell cycle and survival pathways (such as Myc targets, PI3K/AKT/mTOR signalling, p53, mitotic spindle, G2M checkpoints, E2F targets, DNA repair, apoptosis), and metabolism (unfolded protein response, mTORC1 signalling, glycolysis, heme metabolism). In contrast, bradyzoite infection appears to more heavily impact immune pathways (complement, IL6/JAK/STAT3 signalling, inflammatory response, interferon gamma response, interferon alpha response), in addition to some pathways that are also altered by tachyzoites. Others have used microarrays to analyse the expression of bradyzoite-containing host cells and concluded that, when the effects of the stress on the host were taken into account, there was little to no transcriptional differences to tachyzoite infected host cells^70^. However, in this study bradyzoite samples were taken 44 hours post infection as compared to our study that took samples after 7 days. The enhanced sensitivity of RNAseq vs microarray, or residual tachyzoite effects at this early time point may explain the differences in our findings^70^.

It is interesting to note that known targets of GRA16^17^ and the newly identified HCE1/TEEGR effector^23, 24^ are differentially affected between bradyzoite and tachyzoite-infected cells. HCE1/TEEGR targets E2F-dependent genes (including cyclins and NF-kB targets) and these are clearly not affected in bradyzoite infected cells, thus we would predict that this effector is not expressed or exported in latent forms. GRA16 is known to be a key player in host cell cycle regulation by modulating p53 signalling, DNA replication, and DNA repair^17^. In our data, these pathways appear to be regulated to a lower degree in bradyzoites when compared to tachyzoites, in accordance with our observation that GRA16 is not exported in these stages^79^. Furthermore, tachyzoites are known to induce DNA replication but block the G2M transition preventing mitosis in host cells, thus increasing the ploidy of the host cells^80, 81^. These observations complement our data which shows the regulation of both the G2M and mitosis pathways in tachyzoites but not bradyzoites. The differential ability for *Toxoplasma* zoites to control cycle cell pathways could reflect the type of host cell the parasites reside in. Tachyzoites can replicate in rapidly dividing cells, whilst bradyzoites appear to develop predominantly reside in post-mitotic cells, such as neurons, myocytes and cardiomyocytes, and thus may have no need to modulate the cell cycle^82^.

Our work also suggests that MYR1-dependent manipulation of host cells containing bradyzoites allows for their survival during immune attack. The loss of MYR1 causes the expression of many genes to revert to the transcriptomic pattern of uninfected bystander cells, suggesting that bradyzoites actively supress many genes which would normally become active in an inflammatory environment. A good example of this pattern from our data is the IFNγ pathway. The expression of IFNγ related genes globally decreases in bradyzoite-containing cells when compared to uninfected bystanders (presumably by exported IST). Upon the loss of MYR1, the manipulation of IFNγ-mediated gene expression is lost, and bradyzoite-containing cells mimic uninfected cells from the same culture.

A parasite’s ability to interfere with the IFNγ-axis is important because it is the host’s key mechanism in controlling *Toxoplasma* infection *in vivo*^46^. The mechanism by which IFNγ does this *in vivo* is not fully understood. Through *in vitro* analysis of human and murine cells, it’s been suggested that IFNγ induced cell death or nutrient depletion might play a role during acute stages^41, 83^. Our data suggests that during chronic infection IFNγ may also play a role initiating cell death as an anti-microbial tool. We have shown that fibroblasts containing MYR1-knockout bradyzoites undergo cell death upon IFNγ stimulation, whereas this is not the case in tachyzoites stimulated with IFNγ post infection. This suggests that bradyzoite protein effectors may work to block IFNγ mediated death. Furthermore, we show cell death occurs in the absence of parasite egress. It is speculated that in tachyzoite-infected macrophages cell death may occur by pyroptosis initiated by the release of PAMPs following GBP-induced PVM breakdown in human cells^41^, but the type of cell death observed in IFNγ-primed HFFs remains undetermined^84^ and apoptosis could be equally likely. Although type II tachyzoites lacking IST are more susceptible to clearance after and before the addition of IFNγ in murine macrophages^19, 20^, we have shown that cell death upon IFNγ of bradyzoite-containing HFF cells is not solely controlled by IST, suggesting that there could be other parasite effectors in play in chronic stages. Together, these data implies that bradyzoites actively modulate cell pathways that control cell death and that this phenomenon is different between latent and acute forms.

Uncovering the cell death pathway initiated upon IFNγ stimulation and how the parasite inhibits this is important in understanding how the chronic stages persistent. Further studies should look into whether IFNγ-induced cell death occurs in physiologically important bradyzoite reservoirs such as neurons^85^. Unfortunately, the current limitations in *in vitro* bradyzoite-neuron tissue culture models makes this a difficult course. Alternatively, *in vivo* studies could shed light on whether MYR1 is required for bradyzoite persistence, but will require a conditional system as MYR1 knockouts are avirulent in acute stages *in vivo*^28^. Overall, this data suggests that targeting protein export with small drug molecules, or augmenting cell death pathways (as has been used in for clearance of other pathogens)^86^ could be an effective way to treat chronic *Toxoplasma* infection.

## Acknowledgments

We’d like to acknowledge to Niall Geohegan and Cindy Evelyn from WEHI’s Centre of Dynamic Imaging. We’re thankful to Timothy McCulloch and Stacey Jeffrey for assistance with flow cytometry. We are grateful to Anita Koshy for providing us with the Pru-Cre parasite line used and John Boothroyd for providing us with the SRS9 and CST1 antibodies. We’d like to thank Stephen Wilcox from the Ian Potter Centre for Genomics and Personalised Medicine at WEHI for assistance with RNA sequencing and Roberto Bonelli for assistance with R modelling. SS is supported by the Australian Government as a recipient of the Research Training Program Stipend Scholarship. We’re grateful for institutional support from the Victorian State Government Operational Infrastructure Support and the Australian Government NHMRC IRIISS. Finally, we would like to acknowledge The David Winston Turner Endowment Fund for supporting this work.

## Author Contributions

SS designed and performed the experiments, analysed the data, and generated the figures. MJC preformed experiments. LWW produced code to analyse imaging data. ALG analysed RNA sequencing data and generated figures. CJT supervised all work. SS and CJT wrote the manuscript with input from all authors.

## Data availability

The RNA sequencing dataset produced during this study have been deposited in Gene Expression Omnibus (GEO), accession code GSE125122, reviewer code: ahspsaaanfqpjyn. Further data is available from the corresponding author upon reasonable request.

---

**Figure S1, related to Figure 2. Pathways enriched during both tachyzoite and bradyzoite infection** (**A**) Over-representation analysis results for tachyzoite vs uninfected (grey) and vs uninfected DEGs (green) for each Hallmark gene set displayed as the negative Log_10_ of the p-value, dotted line p = 0.05. (**B**) Over-representation analysis results for tachyzoite and bradyzoite shared DEGs (black) for each Hallmark gene set displayed as the negative Log_10_ of the P-value, dotted line p = 0.05. See also Table S2.

**Figure S2, related to Figure 3. Expression and knockout of MYR1** (**A**) Tachyzoites (αGAP45; green) express MYR1 (αHA; white) at the PV and within the PV space. Scale bar = 10μM (**B**) A random 69 bp insertion between 237-238 results in three stop codons (**C**) Transiently transfected GRA24-myc3 (αMyc; green) export into the host nucleus (DAPI; blue) is inhibited in Δ*myr1* tachyzoites (mCherry; red). White arrows indicate location of host nucleus. Scale bar = 10μM. Brightness and contrast was edited on single colour channels prior to merging.

**Figure S3, related to Figure 3. MYR1 dependent DEGs play a significant role in pathways regulated by bradyzoites infection.** (**A**) Over-representation analysis results for the MYR1 dependent DEGs for each Hallmark gene set displayed as the negative Log_10_ of the P-value, dotted line p = 0.05. (**B**) Over-representation analysis results for the MYR1 independent DEGs (black) for each Hallmark gene set displayed as the negative Log_10_ of the P-value, dotted line p = 0.05. (**C**) RPKM values for XIAP protein expression Figure 4 data set. See also Table S5 for gene lists.

**Figure S4, related to Figure 5. Bradyzoites export IST.** (**D**) Quantification of the intensity of nuclear GRA16-HA, GRA24-HA and IST-HA (normalised to PruΔ*ku80* WT controls) of bradyzoites infected host cells from immuno-fluorescence imaging (Figure 6c) (mean ± SD, n = 3 experiments, each experiment analysed on average between 20-40 bradyzoite cysts and 60-200 tachyzoite vacuoles; unpaired, parametric one-way ANOVA test with Dunn’s multiple comparison; ****p<0.0001)

**Figure S5, related to Figure 6. MYR1 knockouts effects on IRF1** (**A**) Western blot indicates IST (αHA) is expressed in WT and Δ*myr1* tachyzoites but is absent in Δ*ist*. αGAP45 is used as a loading control. (**B**) Immuno-fluorescence images of tachyzoite (αGAP45; green) PruΔ*ku80* IST-HA (WT), PruΔ*ku80* IST-HA Δ*myr1*, and PruΔ*ku80* IST-HA Δ*ist* infected cells. WT can export IST-HA (αHA; white) in to the host nucleus (DAPI; blue), whereas there is a loss of export in Δ*myr1* and no expression in Δ*ist* infected cells. Scale bar = 10μM. Brightness and contrast was edited on single colour channels prior to merging. (**C**) (i) PruΔ*ku80* IST-HA (WT), PruΔ*ku80* IST-HA Δ*myr1*, and PruΔ*ku80* IST-HA Δ*ist* bradyzoites (green) were treated with 50ng/ml of human-IFNγ for 15hrs and before being fixed and probed for IST (anti-HA; white) and IRF1 (magenta). There is a range of IRF1 inhibition. (iii) The relationship between the intensity of IRF1 and the intensity of nuclear IST-HA was modelled with a non-linear curve (blue) using natural splines with 3 degrees of freedom. This could significantly explain the relationship between IST-HA and IRFI (p=1.042×10^−12^) (**D**) RHΔ*ku80* IST-HA tachyzoites and PruΔ*ku80* IST-HA parasites, that were either differentiated under alkaline conditions for 7 days (bradyzoites) or grown in D1 for 15 hours (tachyzoites), were treated with 50ng/ml of human-IFNγ for 15hrs. Samples were fixed and probed with αIRF1 and DAPI. Nuclear IRF1 protein intensity levels were measure as a proportion of surrounding uninfected cell intensity for both uninfected and infected cells. There was no difference in IRF1 expression between PruΔ*ku80* tachyzoites and bradyzoites (mean ± SD, n = 3 experiments, p=0.306). However there is a significant difference between PruΔ*ku80* and RHΔ*ku80* tachyzoites (****p<0.0001). Each experiment analysed on average between 20-30 bradyzoite infected host cells and 60-200 tachyzoite vacuoles (unpaired, one-way ANOVA test with Sidak’s multiple comparison test).

**Figure S6, related to Figure 6. RPKM values for protein expression of IFNγ-pSTAT1-GAS induced transcripts from** Figure 4 **data set.** Samples include: uninfected (green), bradyzoite WT-infected (pink) and bystander (orange), bradyzoite Δ*myr1*-infected (indigo) and bystander (aqua) cells.

**Figure S7, related to Figure 7. IFNγ induced cell death of bradyzoite containing HFF cells.** (**A**) PruΔ*ku80* IST-HA (WT), Δ*myr1*, and Δ*ist* tachyzoites were treated with IFNγ for 15 hours. HFFs were washed, trypinised and fixed, before probed with αGAP45. Samples were analysed using FACS (mean ± SD, n = 5 experiments, unpaired, one-way ANOVA test with Sidak’s multiple comparison test; p>0.98) (**B**) Compound 1 (2μM) was added to 7-day-old PruΔ*ku80* IST-HA Δ*myr1* bradyzoites (green) for an hour prior to the IFNγ addition for 15 hours. PI (red) was added for 10 mins before being washed and fixed, white arrows indicated PI positive parasites. (**C**) Compound 1 (2μM) was added to 7-day-old PruΔ*ku80* IST-HA Δ*myr1* bradyzoites for 16 hours prior to the addition of BIPPO or vehicle (DMSO). Samples were trypinised and fixed, FACS analysed, and the proportion of bradyzoite-eGFP^+^ cells compared to untreated samples (mean ± SD, n = 3 experiments, unpaired, one-way ANOVA test with Sidak’s multiple comparison test; p<0.0001, p=0.889).

**Video S1, related to Figure 7. Wildtype bradyzoite lysis during IFNγ stimulation.** Parasites were differentiated for 7 days (bradyzoites express pLDH2-eGFP, green), before the addition of Propidium Iodide (red) and IFNγ and imaged every 14 mins for 15 hours.

**Video S2, related to Figure 7. Δ*myr1* bradyzoite lysis during IFNγ stimulation.** Parasites were differentiated for 7 days (bradyzoites express pLDH2-eGFP, green), before the addition of Propidium Iodide (red) and IFNγ and imaged every 14 mins for 15 hours.

**Video S3, related to Figure 7. Δ*ist* bradyzoite lysis during IFNγ stimulation.** Parasites were differentiated for 7 days (bradyzoites express pLDH2-eGFP, green), before the addition of Propidium Iodide (red) and IFNγ and imaged every 14 mins for 15 hours.

**Table S1. Log RPKM values for the top 100 differentially regulated genes between uninfected host cells and bradyzoite infected host cell related to Figure 1D**.

**Table S2. Table of DEG and their corresponding Hallmark Pathways for tachyzoite specific, bradyzoite specific and shared genes related to Figure 2A and S1.**

**Table S3. Log RPKM values for genes expressed in the E2F, Myc, IFNα and IFNγ Hallmark pathways for tachyzoite (related to Fig. 2B) and bradyzoite (related to Fig. 2C) comparisons.**

**Table S4. Log RPKM values for the top 100 differentially regulated genes between bradyzoite WT-infected and Δmyr1-infected host cells related to Figure 3D**.

**Table S5. Table of DEG and their corresponding Hallmark Pathways for MYR1 dependent (WT and Δ*myr1* specific) genes and MYR1-independent (shared) genes related to Figure 4 and S3.**

**Table S6. Log RPKM values for genes expressed in the IFNγ (related to Fig. 6) Hallmark pathways for MYR1 knockout comparisons.**

